# Kaposi sarcoma herpesvirus vFLIP induces lipid droplet formation in Tert-immortalized microvascular endothelial cells via NF-κB activation

**DOI:** 10.64898/2026.10.22.760464

**Authors:** Blake D. Dodson, Mariah E. Thompson, Ryan W. Tibble, Patrick S. Mitchell, Michael Lagunoff

## Abstract

Kaposi sarcoma herpesvirus (KSHV) is the etiologic agent of Kaposi sarcoma (KS) and preferentially establishes a latent infection state in both the characteristic spindle cells of the KS tumor and cultured endothelial cells. KSHV has been described to broadly alter cellular metabolism during latent infection of endothelial cells. In Tert-immortalized microvascular endothelial (TIME) cells, our lab has shown that latent infection increases the abundance of long chain fatty acids in the cell, as well as formation of lipid droplets, which is dependent on viral gene expression. Here, we show that acyl-CoA synthetase activity, crucial for lipid droplet formation, is essential for cell survival during latent infection of TIME cells. Expression of the major genetic locus expressed during KSHV latency is sufficient to induce lipid droplet formation. Virus mutants lacking expression of the latent viral gene vFLIP, or the expression of the latent viral microRNAs, do not induce lipid droplet formation. Expression of vFLIP in isolation is sufficient to induce lipid droplet formation and sensitivity to ACSL inhibition, but a mutant version of vFLIP with decreased NF-κB signaling capacity is unable to increase lipid droplet formation to the same degree as its wild-type counterpart. Lipid droplet formation is also limited in the context of co-expression with a mutant version of IκBα resistant to degradation. We conclude that vFLIP drives the increase in lipid droplet formation observed during latent KSHV infection, in an NF-κB signaling dependent fashion and that ACSL function is important for the survival of latently infected endothelial cells.

**Importance:** Kaposi’s sarcoma herpesvirus is the etiologic agent of Kaposi’s sarcoma (KS). The main tumor cell of KS, the spindle cell, expresses markers of endothelium. Spindle cells and KSHV infected cultured endothelial cells are both predominantly latently infected. How KSHV alters host endothelial cells to cause KS is an active area of study. During latent infection, KSHV dramatically alters cellular metabolism, including inducing long chain fatty acids and lipid droplets, a storage organelle for fatty acids. This study shows that inhibition of the enzyme necessary for creating lipid droplets decreases survival of KSHV infected endothelial cells and a single KSHV latent gene, vFLIP, is both necessary and sufficient for induction of lipid droplets. Overall, this study identifies lipid droplet formation as an important host process for KSHV latent infection and identifies both the viral and cellular mechanism for the induction of lipid droplets.

## Introduction

Kaposi sarcoma herpesvirus (KSHV) is an oncogenic virus and the etiologic agent of Kaposi sarcoma (KS), as well as the B cell malignancies primary effusion lymphoma and multicentric Castleman’s disease, and the inflammatory disease known as Kaposi sarcoma inflammatory cytokine syndrome^1–5^. KS has a significant impact on global health, as it is one of the most common malignancies in AIDs patients across the world, and one of the most common forms of cancer in countries of eastern, central, and southern Africa where KSHV is endemic^6,7^. It primarily presents as a complex, multifocal tumor of the skin that displays an extensive degree of vascularization, and the characteristic proliferating cell type of these tumors, termed spindle cells, are known to express endothelial cell markers^8,9^.

A member of the gammaherpesvirus family, KSHV has a large dsDNA genome encoding over 90 open reading frames and a biphasic replication scheme consisting of lytic and latent replication cycles^10,11^. Latency is the dominant state observed in spindle cells in the KS tumor, with only a small minority of these cells expressing markers of lytic replication and producing progeny virions^8,12,13^. *In vitro* infection of primary endothelial cells and Tert-immortalized microvascular endothelial (TIME) cells with KSHV also results preferentially in the establishment of latency, with latent and lytic infection observed at similar rates to those observed in spindle cells^14^. Viral latency is established by 48 hours post-infection (hpi) *in vitro* and viral gene expression during latent infection in endothelial cells is largely limited to a single genetic locus, the KSHV latency-associated region (KLAR), though specific genes outside this locus may be expressed sporadically or to very low levels^15–17^. The KLAR consists of 12 microRNA (miRNA) loci and four viral open reading frames, respectively resulting in the production of 25 mature miRNA species and 6 proteins: LANA, vCyclin, vFLIP, Kaposin A, Kaposin B, and Kaposin C^16–19^. During latency, these viral factors manipulate a multitude of cellular pathways to facilitate cell survival and continued proliferation, ensure viral genome maintenance and segregation, and carefully regulate viral latency and lytic reactivation.

Among these latent viral proteins, vFLIP activates constitutive canonical NF-κB signaling by associating with the regulatory subunit of the IKK complex, IKKγ^20^. This interaction triggers the autophosphorylation of IKKβ, allowing it to phosphorylate the NF-κB negative regulator IκBα, which triggers its degradation and allows the nuclear translocation of the NF-κB subunits, RelA (p65) and p50^20,21^. The activated IKK complex also phosphorylates p65 at serine 536 (S536), enhancing its transactivation activity^22^.

Activation of NF-κB is important for the maintenance of viral latency, as NF-κB antagonizes two cellular transcription factors, AP-1 and RBP-Jκ, which drive transactivation of the KSHV lytic switch protein, Rta^23,24^. Additionally, vFLIP expression in endothelial cells has been reported to result in an increase in expression of cyclooxygenase-2 and activation of several MAP kinases, suggesting a broader range of roles for vFLIP in manipulating the cell in addition to those currently appreciated^25^.

Treatment of KS is currently limited to standard cancer chemotherapy, as therapeutically targeting the latent virus has proven challenging due to the small number of viral targets expressed, and a true cure for the disease remains elusive^13^. As such, there is great interest in identifying changes in cell state induced and rendered essential for cell survival during latent infection, including changes in cellular metabolism. Our previous metabolomics study revealed lipid metabolism to be significantly altered during latent infection of TIME cells by KSHV, as two-thirds of detectable long-chain fatty acid species were increased in abundance in infected TIME cells relative to mock infected controls at 48 and/or 96 hpi, while none were decreased^26^. Inhibitors of fatty acid synthesis led to significantly increased cell death in KSHV latently infected endothelial cells but not in uninfected cells, indicating that fatty acid synthesis is required for the survival of KSHV latency^26^.

High levels of free fatty acids pose a danger to the cell, as they can cause oxidative and ER stress in a metabolic phenotype known as lipotoxicity^27,28^. Cells may alleviate these stressors by converting free fatty acids into acyl-CoA thioesters via the activity of a family of enzymes called acyl-CoA synthetases (ACSLs) and packaging these acyl-CoAs into neutral lipid species, such as triglycerides and cholesterol esters^27^. These neutral lipids are then stored in cytoplasmic lipid droplets until the cell requires them for energy production, membrane synthesis, or synthesis of bioactive lipid signaling molecules^27^. Our previous investigation into KSHV’s impact on TIME cell metabolism also observed an increase in cytoplasmic lipid droplet formation during latent infection of TIME cells^26^. Furthermore, this induction of lipid droplet formation was observed to be dependent upon viral gene expression, as infection with UV-irradiated virions did not yield the same phenotype^26^. Although this suggests a viral factor is responsible for the observed increase in lipid accumulation in endothelial cells during latency, the viral mechanism by which latent KSHV induces lipid droplet formation remains uncharacterized, as does the importance of this induction of lipid droplet formation to cell survival during latent infection.

Here, we found that ACSL inhibition by Triacsin C kills latently infected TIME cells and that the KSHV latent protein vFLIP is essential for induction of lipid droplets during infection of TIME cells. vFLIP is individually sufficient to induce lipid droplet formation in TIME cells, an effect achieved by virtue of its activation of NF-κB. These studies provide a more detailed understanding of how KSHV alters cellular lipid metabolism and suggest that lipid accumulation could be an intriguing target for further therapeutic lines of inquiry.

## Results

### Acyl-CoA synthetase inhibitor Triacsin C treatment kills KSHV latently infected TIME cells

Previous investigations into the alteration of endothelial cell metabolism during latent KSHV infection revealed that latent infection triggers lipid droplet formation in TIME cells^26^. To determine whether this process is essential for cell survival during latent infection, we aimed to pharmacologically inhibit the process of lipid droplet formation using Triacsin C, the only commercially available broad-spectrum inhibitor of the acyl-CoA synthetase (ACSL) family, which has been well established to decrease lipid droplet formation^33–37^. Treatment with 5 μM Triacsin C for 48 hours did not have a significant impact on cell viability in mock infected TIME cells relative to DMSO-treated control cells but led to a significant decrease in cell viability in TIME cells latently infected with KSHV, as assessed by trypan blue exclusion assay (**Figure 1A**).

**Figure 1:**
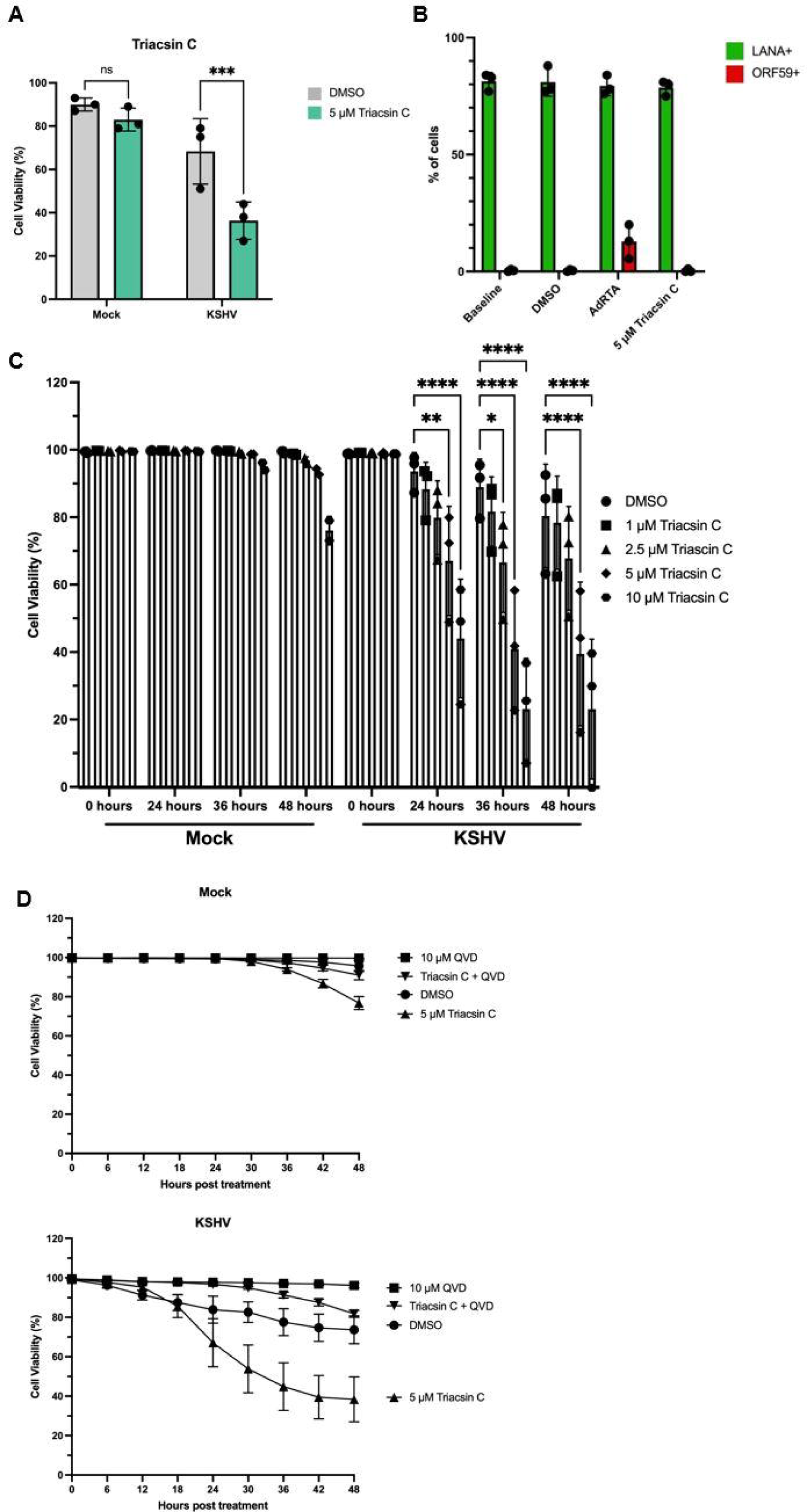
Acyl-CoA synthetase inhibitor Triacsin C treatment kills KSHV latently infected TIME cells. (A) Cell viability measured by trypan blue exclusion assay. Mock and KSHV infected TIME cells were treated with 0.3% DMSO or 5 uM Triacsin C at 48 hpi. 48 hours following drug treatment, cell viability was assessed. (B) Percentage of infected TIME cells displaying staining for LANA (green) and ORF59 (red) following immunofluorescence. KSHV infected TIME cells were pooled at 48 hpi and split into three groups. A sample of cells was taken at 48 hpi and fixed to establish baseline levels of latent and lytic infection prior to drug treatment/adenovirus transduction. Two of the groups were then treated with 0.3% DMSO or 5 uM Triacsin C, while the third was transduced with a gutted adenovirus expressing the KSHV lytic switch protein (AdRTA). At 24 hours post treatment/transduction, cells were fixed and latent and lytic infection were assessed by immunofluorescence. Data shown are mean +/-SD, n=3. 2-way ANOVA with Tukey’s test was used. (C) Assessment of dose response to Triacsin C treatment in mock and KSHV infected TIME cells beginning at 48 hours post-infection. Cell viability was assessed over time using the fluorescent green total cell dye SYTO 24 Green and the fluorescent red dead cell dye propidium iodide in combination with a CellCyte live cell imager. Cell viability measurements are shown at 0, 24, 36, and 48 hours post-drug treatment and dye staining. Data shown is mean +/- SD, n=3. Two-way ANOVA with Dunnett’s multiple comparisons test was used. (D) Mock (top) and KSHV latently infected (bottom) cells treated with Triacsin C in isolation or in combination with the pan-caspase inhibitor Q-VD-OPh (QVD). Cell viability was assessed as in (C); data shown is mean +/- SD, n=3.

Rates of latent infection and lytic reactivation were assessed via immunofluorescence for LANA, the master regulator of viral latency, and ORF59, a lytic replication marker, respectively. These measures were assessed from a pool of latently infected cells pre-drug treatment at 48 hpi (baseline), as well as 24 hours following treatment with DMSO, treatment with Triacsin C, or transduction of RTA into latently infected cells using a gutted adenovirus (AdRTA)^32^. No lytic reactivation was observed in the population of latently infected cells at baseline or following 24 hours of treatment with DMSO or Triacsin C (**Figure 1B**). Importantly, these cells showed no inherent defect in lytic reactivation, as transduction of RTA was able to induce lytic reactivation in each experimental replicate (**Figure 1B**). Rates of latent infection were similar across all groups (**Figure 1B**).

Further examination of the sensitivity of KSHV latently infected TIME cells via live cell imaging revealed dose-and time-dependent effects of Triacsin C treatment on cell viability during latency with a dose-dependent decrease in cell viability observed at 24 hours post Triacsin C treatment and increasing at the 36- and 48-hour timepoints (**Figure 1C**). Co-treatment of Triacsin C treated cells with the pan-caspase inhibitor Q-VD-OPh (QVD) was able to rescue this decrease in cell viability (**Figure 1D**). Together, these data suggest that ACSL inhibition by Triacsin C treatment is detrimental to the survival of TIME cells during KSHV latent infection, and that this survival requirement is a function of the latent infection state of the cell. The violation of this survival condition leads to programmed cell death in a caspase-dependent manner, rather than occurring as a result of lytic reactivation or ferroptotic cell death.

### Expression of the KSHV major latent locus is sufficient to drive lipid droplet formation in TIME cells

Previous studies revealed that the observed increases in lipid droplet formation during KSHV latent infection required KSHV gene expression, as cells infected with UV-irradiated KSHV virions did not display increased lipid droplet formation compared to mock infected controls^26^. The major genetic locus expressed by the virus during latent infection of endothelial cells, termed the KSHV latency-associated region (KLAR), is composed of 6 viral genes: LANA, vCyclin, vFLIP, Kaposin A, Kaposin B, and Kaposin C, as well as 12 latent miRNA loci^18^. To determine whether expression of the KLAR is sufficient to induce lipid droplet formation in TIME cells, a gutted adenovirus (AdKLAR) was used to transduce the latent locus under the native promoter without expression of any adenoviral genes (**Figure 2A**)^32^. LANA expression was confirmed in AdKLAR transduced cells via RT-qPCR (**Supplementary Figure 1A and 1B**). As a control for adenoviral attachment and entry, an equivalent number of viral particles of a similar gutted adenovirus expressing only Rta of KSHV and no other adenoviral or KSHV genes, AdRTA, was also used. Lipid droplet formation was assessed as previously described, using the fluorescent neutral lipid dye Nile Red, and assessing fluorescence via flow cytometry^26^. As previously observed, KSHV latently infected TIME cells display a rightward shift in Nile Red fluorescence, and thus a significant increase in lipid droplet formation compared to their mock infected counterparts at 48 hpi (**Figure 2B and 2C**). Cells transduced with AdKLAR displayed a similar rightward shift in Nile Red fluorescence, indicating a similar increase in lipid droplet formation, compared to cells transduced with AdRTA at 48 hours post-transduction (**Figure 2B and 2C**). Therefore, expression of the major KSHV genetic locus expressed during latent infection is individually sufficient to induce lipid droplet formation.

**Figure 2:**
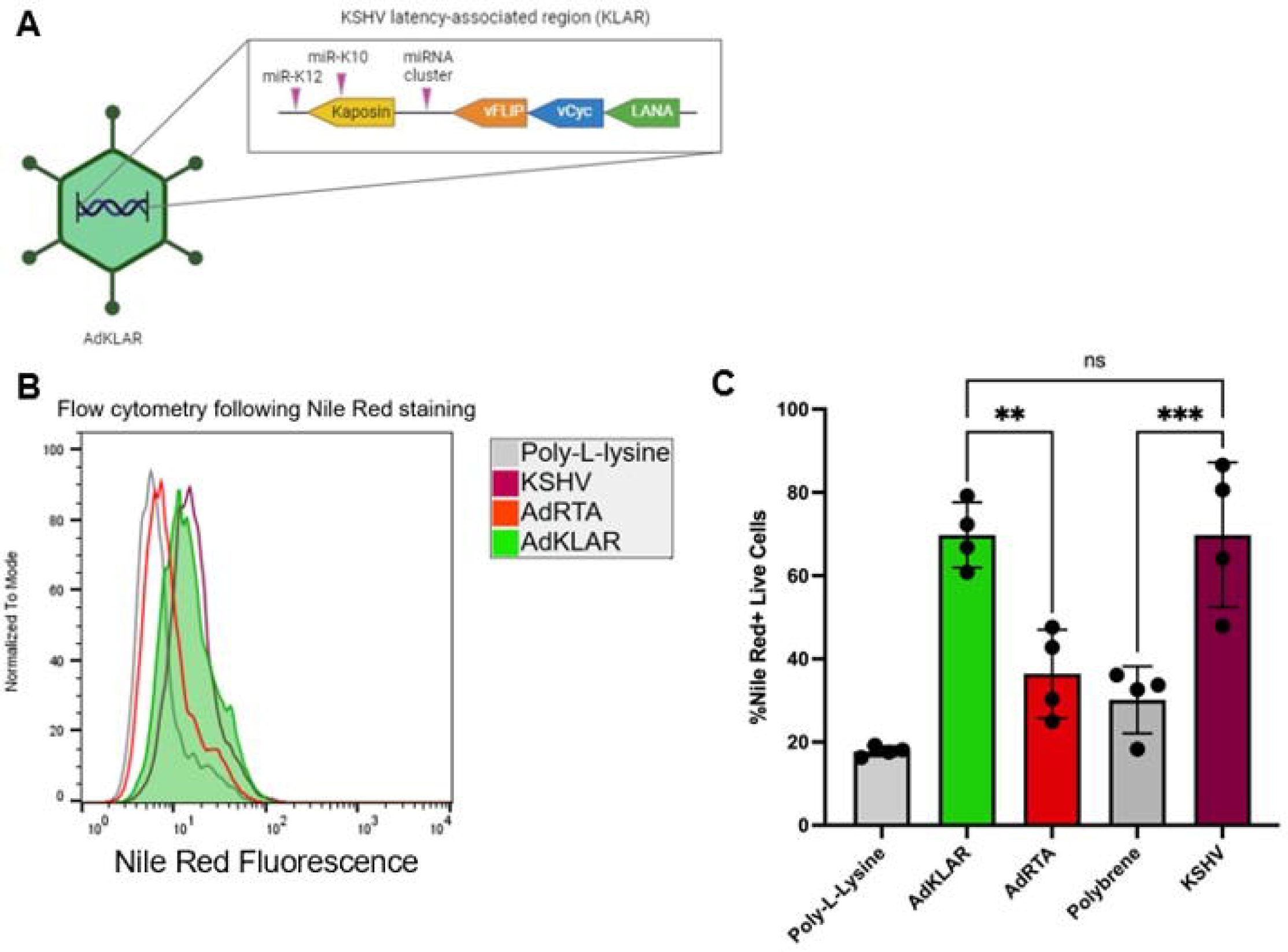
Expression of the KSHV major latent locus is sufficient to drive lipid droplet formation in TIME cells. (A) The AdKLAR gutted adenovirus. (B) Flow curves of populations of cells treated with poly-L-lysine (gray), infected with KSHV (maroon), transduced with AdRTA (red), or transduced with AdKLAR (green), then fixed and stained with Nile Red at 48 hours post-infection/transduction. Dead cells and debris have been gated out. (C) Quantification of flow curves generated by setting a fluorescence range gate determined by the poly-L-lysine-treated, negative control cell population. Data shown is mean +/- SD, n=4. Two-way ANOVA with Tukey’s test was used.

### KSHV latent protein vFLIP and the microRNA loci are essential to the induction of lipid droplet formation observed during KSHV latent infection of TIME cells

We obtained deletion virus mutants for the KSHV latent factors vCyc, vFLIP, Kaposins and the miRNA locus^29,30,38^. These deletion viruses were used to determine which of the viral latent factors encoded within the KLAR are essential for the induction of lipid droplets. As the deletion viruses lacking expression of vCyclin (ΔvCyc), vFLIP (ΔvFLIP), or all 12 of the latent miRNA loci (ΔmiRNAs) express GFP, the red fluorescent neutral lipid dye LipidSpot 610 was used in place of Nile Red. TIME cells latently infected with KSHV display a rightward shift in LipidSpot 610 fluorescence relative to their mock infected counterparts, as do cells latently infected with ΔvCyc (**Figure 3A**). However, TIME cells latently infected with ΔvFLIP or ΔmiRNAs do not display the same shift in fluorescence at 48 hpi, instead more closely resembling mock infected cells than cells latently infected with KSHV (**Figure 3B, 3C, and 3D**).

**Figure 3:**
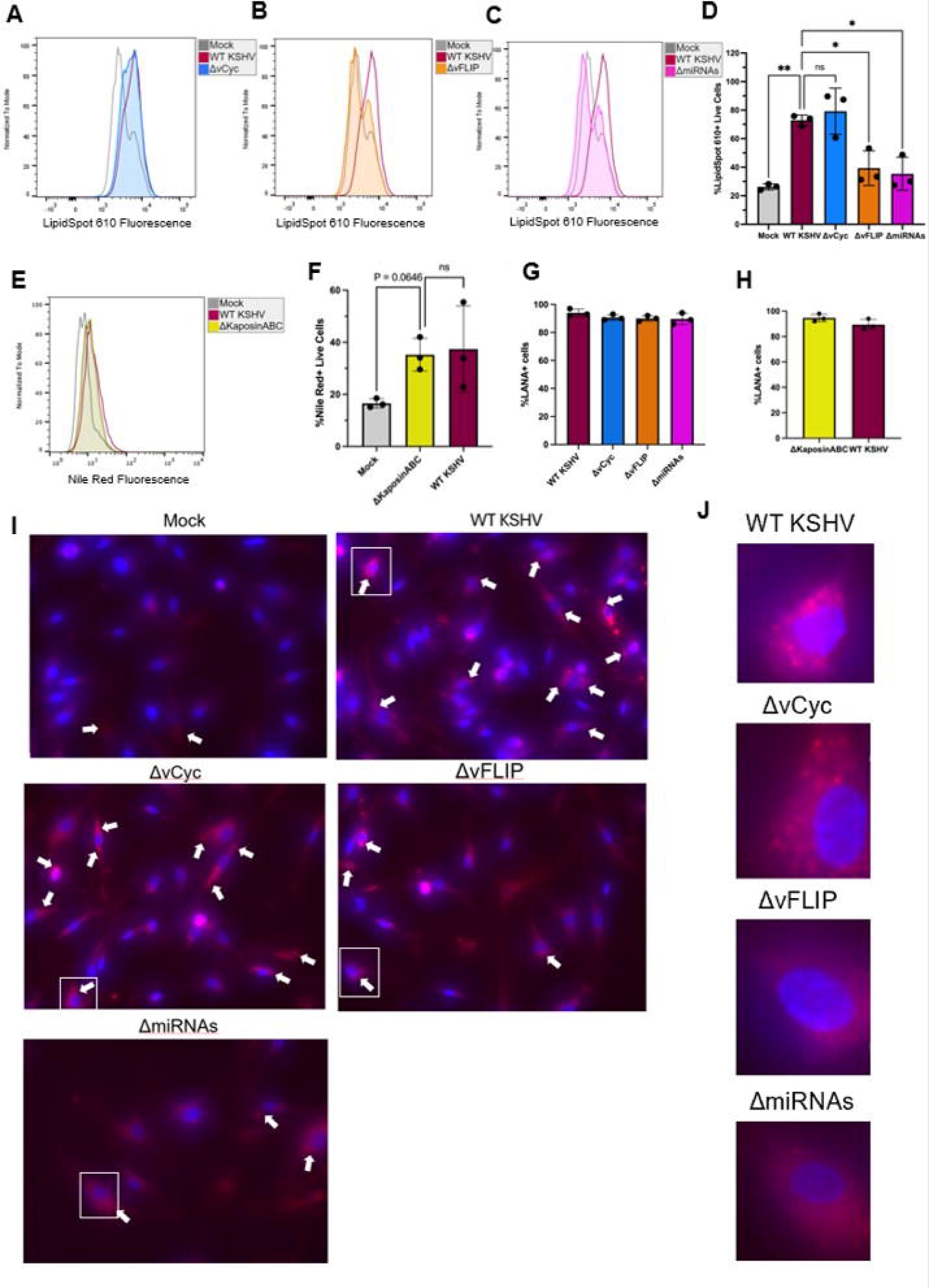
KSHV latent protein vFLIP and latent microRNAs are essential to the induction of lipid droplet formation observed during KSHV latent infection of TIME cells. (A, B, C) Flow curves of live TIME cells infected with KSHV.219 ΔvCyclin (A, blue), ΔvFLIP (B, orange), or ΔmiRNAs (C, pink) compared to mock infected TIME cells (gray) and WT KSHV.219 infected TIME cells (maroon) at 48 hpi. Flow curves are shown from a single experimental replicate. (D) Quantification of A, B, and C. Data shown are mean +/- SD, n=3. Two-way ANOVA with Tukey’s test was used. (E) Flow curves of live TIME cells mock-infected (gray), infected with WT KSHV.219 (maroon), or infected with KSHV.219 ΔKaposinABC (yellow), fixed and stained with Nile Red at 48 hpi. (F) Quantification of E. Data shown are mean +/- SD, n=3. Two-way ANOVA with Tukey’s test was used. (G and H) Percentage of cells staining positive for LANA via immunofluorescence in each experimental replicate of D and F, respectively, at 48 hpi. (I) Fluorescence microscopy of cells latently infected with the indicated virus, fixed and stained with LipidSpot 610 at 48 hours post-infection. Arrows denote cells with visible lipid droplet formation. (J) Fluorescence microscopy images of selected cells from I, denoted by the white boxes in I.

The virus containing a deletion of the Kaposin locus expresses RFP, and therefore Nile Red, which stains lipid droplets green, was used to assess lipid droplet formation. Both WT KSHV and KSHV lacking expression of kaposins A, B, and C (ΔKaposinABC) exhibited a similar increase in Nile Red fluorescence and therefore lipid droplet formation at 48 hpi, indicating that the kaposins are dispensable for the induction of lipid droplet formation during latent infection (**Figure 3E and 3F**).

Lack of expression of the various latent viral factors did not preclude the establishment of latent infection in >85% of the cell population in any of the infections described here, as assessed by immunofluorescence for LANA at 48 hpi (**Figure 3G and 3H**). Fluorescence microscopy conducted using latently infected cells fixed and stained with LipidSpot 610 at 48 hpi corroborated the results observed via flow cytometry and confirmed staining of punctate, cytoplasmic lipid droplets (**Figure 3I and 3J**). Thus, we conclude that vFLIP and the presence of the viral miRNA loci are essential for the induction of lipid droplet formation during latent infection.

### vFLIP is sufficient to induce lipid droplet formation and sensitivity to Triacsin C

To determine whether vFLIP is also sufficient to induce lipid droplet formation in TIME cells, the vFLIP open reading frame was cloned into the lentiviral expression vector pLVX. TIME cells were transduced with pLVX GFP and pLVX vFLIP via lentivirus and successfully transduced cells were selected by puromycin treatment from 24 to 72 hours post-transduction. 48 hours following selection was sufficient to allow LipidSpot 610 staining levels in GFP transduced cells to return to those observed in mock infected cells following 48 hours in culture. At 120 hours post-transduction, cells expressing vFLIP had significantly increased levels of LipidSpot 610 fluorescence relative to those expressing GFP (**Figure 4A and 4B**). This result was further corroborated by fluorescence microscopy using LipidSpot 610, in which LipidSpot 610 staining of lipid droplets is visible (**Figure 4C and 4D**).

**Figure 4:**
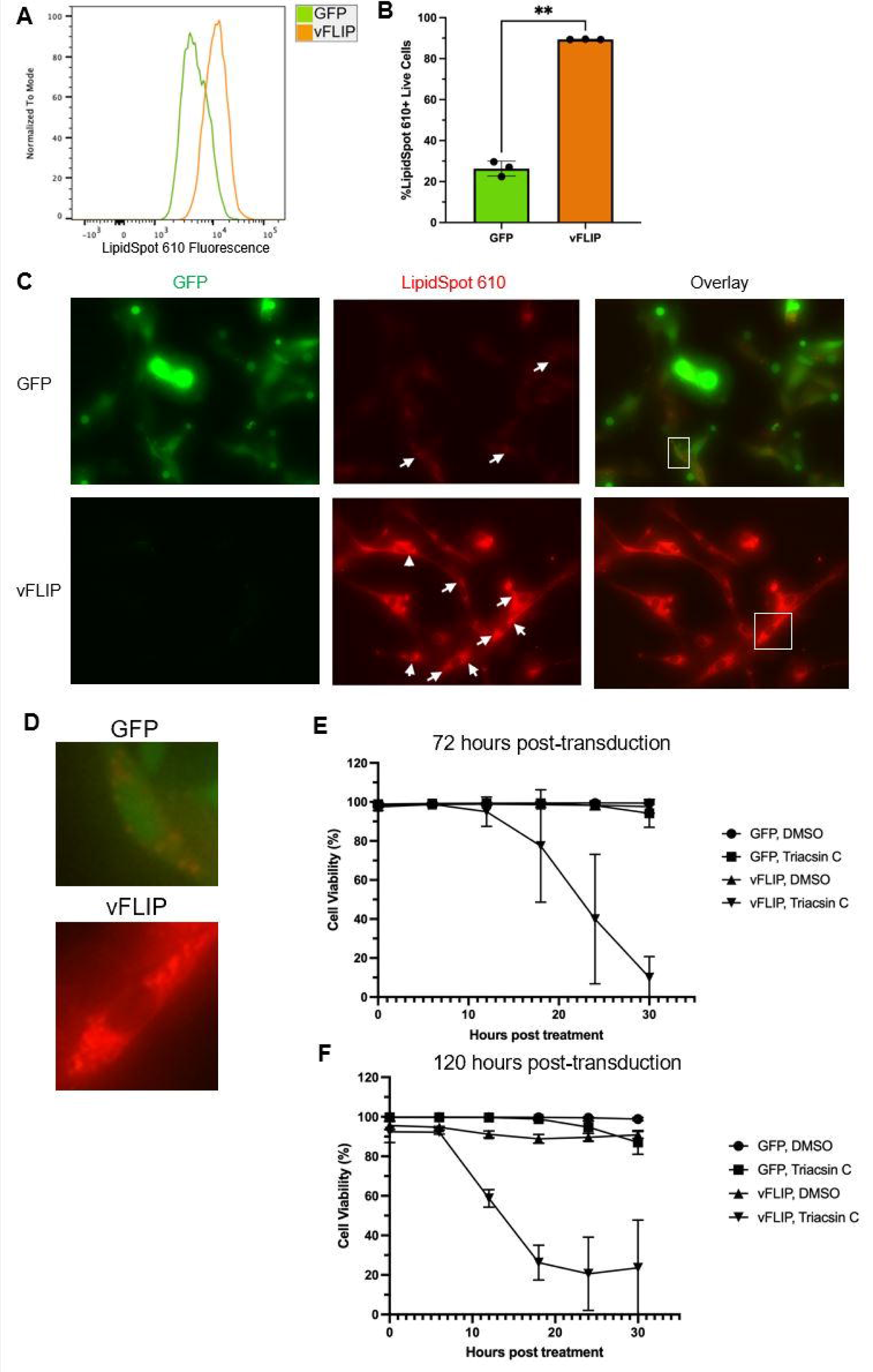
vFLIP is sufficient to induce lipid droplet formation and sensitivity to Triacsin C. (A) Flow curves of LipidSpot 610 staining in live cells overexpressing GFP (green) or vFLIP (orange) at 5 days post transduction. (B) Quantification of A. Data shown are mean +/- SD, n=3. Paired t test was used. (C) Fluorescence microscopy of cells overexpressing GFP (top) and vFLIP (bottom) fixed and stained with LipidSpot 610 at 120 hours post-transduction. White arrows point to visible, punctate cytoplasmic staining of lipid droplets. Identical settings were used to capture both images at a total magnification of 40X. (D) Fluorescence microscopy images of selected cells from C, denoted by white boxes in C. (E) Cell viability assessed over time using fluorescent dead and total cell dyes in combination with a CellCyte live cell imager. Cells were transduced with lentiviruses carrying overexpression vectors for GFP and vFLIP, selected for successfully transduced cells, and treated with 0.1% DMSO or 5 uM Triacsin C at the conclusion of selection at 72 hours post-transduction (top), or provided an additional 48 hours to recover, as in the lipid droplet assessment (bottom). Data shown are mean +/- SD, n=3.

We next wanted to determine if cells expressing vFLIP displayed sensitization to the ACSL inhibitor, Triacsin C. Cells were transduced with pLVX GFP and pLVX vFLIP lentiviruses and were treated with 5 μM Triacsin C beginning at 72 and 120 hours post-transduction. These timepoints were chosen to determine sensitivity to ACSL inhibition prior to the timepoint at which lipid droplets were previously measured and at the time of measurement, respectively. Cell viability was assessed over the next 30 hours via live cell imaging using fluorescent total and dead cell dyes. In cells treated at 72 hours post-transduction, Triacsin C had no significant impact on TIME cells expressing GFP, while nearly 100% of those expressing vFLIP died within 30 hours (**Figure 4E**). In those treated at 120 hours post-transduction, over 70% of cells expressing vFLIP died by 18 hours post-drug treatment (**Figure 4F**). Expression of vFLIP was verified at both timepoints by RT-qPCR (**Supplementary Figure 1C**). These data suggest that vFLIP expression is individually sufficient to drive lipid droplet formation in TIME cells, as well as sufficient to alter cell metabolism to render TIME cells sensitive to Triacsin C treatment.

### vFLIP’s NF-κB signaling capacity is essential to its induction of lipid droplet formation

The binding interface enabling vFLIP to bind IKKγ and trigger NF-κB signaling has been characterized, and it has been reported that the single amino acid mutant vFLIP R12E displays a decrease in NF-κB signaling relative to the WT vFLIP^20^. This mutation was made in the pLVX vFLIP expression construct and the mutant was tested for lipid droplet formation. Expression of vFLIP R12E was assessed via RT-qPCR in four out of the five experimental replicates conducted and, in each case, vFLIP R12E displayed equivalent or higher expression than its WT counterpart (**Figure 5A**). Expression of vFLIP R12E in TIME cells resulted in significantly lower levels of lipid droplets (35% decrease) than those observed in cells expressing the WT vFLIP, but lipid droplet formation was still higher than that observed in cells transduced with pLVX GFP (**Figure 5B and 5C**). To further investigate this intermediate phenotype, as well as the noticeable variability observed between experimental replicates, we assessed vFLIP R12E’s deficiency in driving NF-κB signaling relative to the WT vFLIP using a luciferase reporter assay conducted in HEK 293Ts. There was a consistent, but not statistically significant, defect in NF-κB signaling in cells transfected with pLVX vFLIP R12E of approximately 20% less relative reporter activity compared to those transfected with pLVX vFLIP in equivalent amounts, explaining the intermediate lipid droplet decrease seen with this mutant (**Figure 5D**). Western blotting for phosphorylation of p65 serine 536 (S536) in TIME cells aligned with the results of the luciferase assay, as vFLIP R12E induced a consistently decreased level of p65 phosphorylation similar in magnitude to that observed in the luciferase assay, albeit not statistically significantly (**Figure 5E and 5F**).

**Figure 5:**
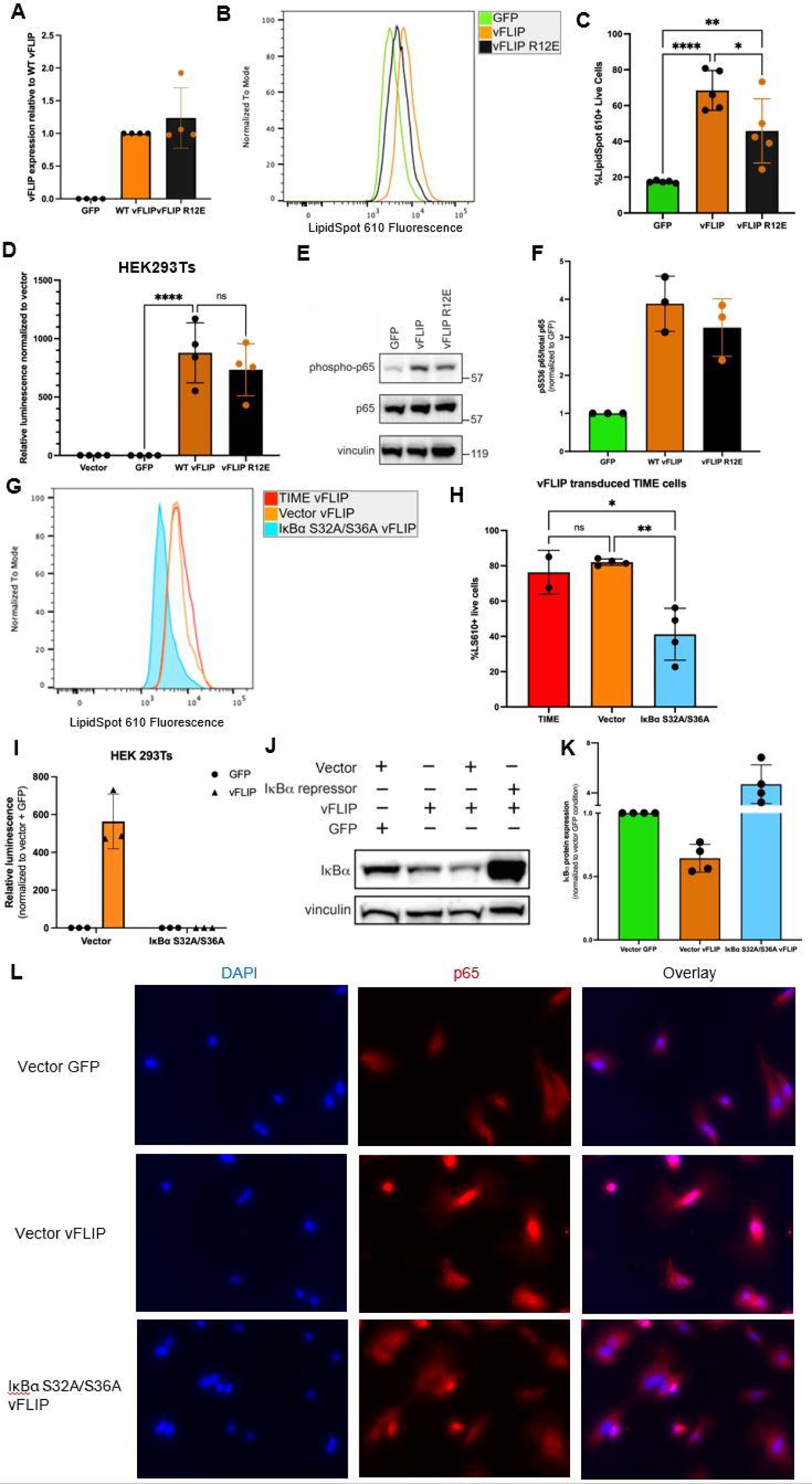
vFLIP’s NF-κB signaling capacity is essential to its induction of lipid droplet formation. (A) Measurement of vFLIP expression relative to cells transduced with WT vFLIP via qRT-PCR. Relative expression was calculated using the 2^-ΔΔCt method with GAPDH or tubulin used to normalize to total RNA. Data shown are mean +/- SD, n=4. (B) Flow curves of live TIME cells overexpressing GFP (green), WT vFLIP (orange), or vFLIP R12E (black), fixed and stained with LipidSpot610 at 120 hours post-transduction. (C) Quantification of B. Data shown are mean +/- SD, n=5. Two-way ANOVA with Tukey’s test was used. (D) Measurement of NF-kB activation via a luciferase reporter assay in 293T cells leveraging firefly luciferase under the NF-kB promoter and NanoLuc luciferase under a constitutive TK promoter. Relative luminescence was calculated by first calculating the ratio of firefly luciferase luminescence to NanoLuc luciferase luminescence for each well, then dividing the firefly:NanoLuc ratio for GFP, WT vFLIP, and vFLIP R12E by that for the well transfected with empty vector. Data shown are mean +/- SD, n=4. Two-way ANOVA with Tukey’s test was used. (E) Representative western blot for phosphorylated p65 and p65, with vinculin as a loading control, for lysates obtained from TIME cells at 120 hours post-transduction with lentiviruses expressing GFP, WT vFLIP, or vFLIP R12E. (F) Quantification of phosphorylated p65:total p65 ratio in western blots as shown in E using ImageJ. n=3. (G) Representative flow curve of unmanipulated TIME cells (red), TIME cells expressing empty vector (orange), or TIME cells expressing IκBα S32A/S36A (blue) transduced with a lentiviral vector expressing vFLIP and fixed and stained with LipidSpot 610 at 120 hours post-transduction. (H) Quantification of G, with the LS610 fluorescence range gate based on the vector vFLIP cell population. Data shown are mean +/- SD, n=4. Two-way ANOVA with Tukey’s test was used. (I) Measurement of NF-kB signaling using the same luciferase assay as that in D. n=3. (J) Assessment of IκBα expression and degradation via western blot with vinculin as a loading control, for lysates obtained from TIME cells expressing the indicated factors at 120 hours post-transduction with lentivirus expressing GFP or vFLIP. (K) Quantification of J using ImageJ. n=4. (L) Observation of p65 nuclear translocation via immunofluorescence microscopy for p65 (red) counterstained with DAPI (blue) at 120 hours post-transduction of cells expressing empty vector and GFP (top), empty vector and vFLIP (middle) or IκBα S32A/S36A and vFLIP (bottom). Nuclei with p65 localized within appear magenta in the overlay panels.

To more robustly suppress vFLIP’s activation of NF-κB, we generated TIME cell lines stably expressing an empty vector or IκBα S32A/S36A, a mutant form of IκBα resistant to phosphorylation by IKK and thus resistant to degradation^39^. These cell lines were transduced with a lentivirus expressing vFLIP, and at 120 hours post-transduction the empty vector control cells expressing vFLIP displayed a similar level of lipid droplet formation to unmanipulated TIME cells expressing vFLIP (**Figure 5G and 5H**). In contrast, the cells expressing IκBα S32A/S36A transduced with vFLIP displayed significantly lower levels of lipid droplet formation (**Figure 5G and 5H**). The expression of empty vector or IκBα S32A/S36A had no impact on lipid droplet formation in cells transduced with the same lentiviral vector solely expressing GFP (**Supplementary Figure 2A and 2B**). The capacity of IκBα S32A/S36A to suppress vFLIP-mediated NF-κB signaling was tested in a luciferase assay in HEK 293Ts, where it demonstrated much more effective inhibition of signaling than vFLIP R12E, returning luciferase expression to near baseline levels (**Figure 5I**). IκBα degradation in TIME cells was assessed via western blot, and vFLIP expression led to a consistent decrease in IκBα in the unmanipulated and empty vector control cells, but not in cells expressing IκBα S32A/S36A, as would be expected given the exogenous expression of IκBα in these cells (**Figure 5J and 5K**). Translocation of p65 to the nucleus following its activation and release from cytoplasmic IκBα was assessed via immunofluorescence microscopy. Empty vector control cells expressing vFLIP displayed visible p65 translocation, whereas those expressing GFP did not (**Figure 5L**). p65 nuclear localization was largely limited in cells expressing IκBα S32A/S36A and vFLIP (**Figure 5L**). Expression of IκBα S32A/S36A did not decrease phosphorylation of p65 induced by vFLIP expression, but this has been observed previously and is not surprising considering that residue S536 is a part of the flexible C-terminal transactivation domain, and not the Rel homology region, where IκBα is known to bind^22,40–42^ (**Supplementary Figure 2C**). Thus, expression of IκBα S32A/S36A resulted in a greater inhibition of vFLIP’s activation of NF-κB than did the introduction of the R12E mutation and a greater decrease in lipid droplet accumulation. These data clearly indicate that the ability of vFLIP to activate NF-κB signaling is essential to its induction of lipid droplet formation.

## Discussion

Inhibition of ACSLs by Triacsin C triggers programmed cell death in TIME cells during latent infection by KSHV. However, ACSL function is not essential for the maintenance of latency in these cells, as treatment with Triacsin C does not trigger lytic reactivation preceding cell death. Therefore, ACSL activity is required during latency due to one or more alterations in the cell causing it to become reliant on these enzymes. Triacsin C may kill latently infected cells due to the ACSL family’s key role in neutral lipid synthesis, which allows free fatty acids to be safely stored in cytoplasmic lipid droplets until they are needed by the cell, but the exact nature of this requirement remains unknown^27^. Since long chain fatty acid species are increased in abundance during latent infection and high levels of free fatty acids are deleterious to cell survival, ACSLs may allow latently infected TIME cells to avoid lipotoxicity^26–28^. Indeed, ACSL inhibition has been shown to trigger cell signaling pathways implicated in lipotoxicity and leading to apoptosis, as Triacsin C treatment of HEK 293 cells has been reported to trigger the IRE1 and PERK arms of the unfolded protein response^28,35^. This would be consistent with our observation that Triacsin C induces caspase-dependent programmed cell death in latently infected TIME cells.

ACSLs are also important for activating free fatty acids for the synthesis of other lipid species, such as phospholipids, so the possibility that this broader function is required during latent infection cannot be ruled out. However, in our hands, latently infected TIME cells do not outpace their mock infected counterparts in proliferation, limiting the idea that ACSLs would be essential for cell survival to supply membrane components and facilitate an increased rate of cell growth and division. Recently, ACSL activity has also been linked to scavenging of exogenous fatty acids, raising the possibility that inhibiting ACSL interferes with fatty acid uptake during latent infection, although the rationale for import of exogenous fatty acids being specifically crucial for cell survival remains unclear^43^.

Lipid droplets are known to accumulate to a higher degree in TIME cells latently infected with KSHV than in mock infected TIME cells, and this accumulation is dependent on viral gene expression^26^. The work reported herein provides a clearer picture of how KSHV mechanistically alters these metabolic pathways. Expression of the KSHV major latent locus alone in TIME cells results in an induction of lipid droplet formation similar to that observed in latently infected cells. The latent viral protein vFLIP is essential for KSHV’s induction of lipid droplet formation, as TIME cells infected with a mutant KSHV lacking expression of vFLIP do not have an increased level of lipid droplets and more closely resemble mock infected cells. Expression of vFLIP is independently sufficient to induce lipid droplet formation, and its expression is likewise sufficient to sensitize TIME cells to the ACSL inhibitor Triacsin C, further connecting ACSL activity to lipid droplet formation in KSHV infected cells.

While vFLIP contains DED domains, it is known to activate NF-κB to prevent cell death and maintain viral latency^20,23,24^. We found that a single amino acid mutation in vFLIP previously described to decrease the protein’s signaling through NF-κB also decreases the formation of lipid droplets observed^20^. It is notable that cells expressing this vFLIP mutant, vFLIP R12E, display an intermediary phenotype between the control GFP and the WT vFLIP expressing cells. This is not altogether surprising considering that in our hands, the vFLIP R12E mutant only reduces NF-κB signaling by ∼20% in the context of expression under the hCMV promoter, whether measured in HEK 293Ts by luciferase assay or in TIME cells by western blot for p65 phosphorylation. By generating TIME cells stably expressing a degradation-resistant IκBα mutant, IκBα S32A/S36A, we were able to achieve more robust suppression of vFLIP’s activation of NF-κB and, as a result, observed a more robust decrease in levels of lipid droplet formation induced by vFLIP. While we cannot formally rule out vFLIP’s other known functions as having an impact on cytoplasmic lipid droplets, by interfering in vFLIP’s activation of NF-κB via two distinct mechanisms and observing defects in lipid droplet accumulation that track closely with the magnitude of NF-κB suppression in both cases, we have identified compelling evidence that vFLIP’s capacity to induce lipid droplet formation is dependent on its activation of NF-κB.

NF-κB has been implicated in lipid accumulation in endothelial cells in the atherosclerosis field as well. NF-κB activation in primary human umbilical vein endothelial cells (HUVECs) can increase expression of SREBP2, a transcription factor known to drive lipogenesis and lipid accumulation, lending credence to the idea that vFLIP could drive lipid droplet formation through NF-kB^44^. Additionally, vFLIP expression in human microvascular endothelial cells (HMVECs) has been shown to result in activation of several MAPKs, and notably Akt^25^. Signaling through the PI3K-Akt axis has been shown to drive lipid accumulation in a diverse array of cell types, ranging from human prostate cancer to *Drosophila melanogaster* nurse cells^45–47^. While the initial report that vFLIP expression in HMVECs did not specifically show that activation of Akt was mediated by vFLIP’s activation of NF-κB, a signaling axis consisting of these three factors could provide a further mechanistic explanation for how vFLIP induces lipid droplet formation in TIME cells^25^.

The viral miRNAs also appear essential for the observed increase in lipid droplets, as a deletion virus mutant lacking expression of all the latent viral miRNAs is also deficient in inducing lipid droplet formation. Similarly to vFLIP, the latent viral miRNA miR-K1 has also been shown to increase NF-κB signaling, which it accomplishes by targeting the 3’ UTR of the IκBα transcript^48^. This suggests that the viral miRNAs may act on lipid metabolism via NF-κB signaling as well. Therefore, both vFLIP and the miRNAs could be required to reach a threshold level of NF-κB activation required to induce lipid droplet formation during latent infection. Preliminary studies in our lab using primary human dermal microvascular lymphatic endothelial cells (HMVEC-dly) have indicated that, while KSHV latent infection strongly induces lipid droplet formation, vFLIP is likely not individually essential for this metabolic process to occur (data not shown). We speculate that the level of NF-κB signaling required to drive lipid droplet formation may vary between cell types. In such a model, it is possible that TIME cells require a higher degree of NF-κB activation to accumulate lipid droplets, rendering both vFLIP and the miRNAs essential for inducing lipid droplet production, whereas this threshold is lower in HMVEC-dlys and as such, a lack of expression of either vFLIP or the miRNAs in isolation is insufficient to return lipid droplet formation to baseline, mock infected levels.

In summary, we have characterized a novel metabolic requirement for TIME cell survival during latent infection and characterized the viral mechanism by which latent KSHV induces lipid accumulation in these cells. Triacsin C treatment induces programmed cell death in latently infected TIME cells, possibly due to the ACSL family’s role in lipid droplet formation and the avoidance of apoptosis triggered by lipotoxicity. KSHV vFLIP is necessary and sufficient to induce lipid droplet formation in TIME cells due to its ability to activate NF-κB, and vFLIP is also sufficient to induce sensitivity to ACSL inhibition. Future studies will examine how latent KSHV mechanistically alters cellular lipid metabolism, and how we might target these metabolic alterations to kill the latently infected cells of the KS tumor with greater precision and specificity.

## Materials and Methods

### Cell lines and tissue culture conditions

TIME cells were cultured in endothelial cell basal medium-2 (EBM-2) (CC-3156) supplemented with Lonza EGM-2 MV Microvascular Endothelial SingleQuots Kit (CC-4147) at 37℃ and 5% CO_2_. HEK 293Ts and iSLK.219s were cultured in Dulbecco’s Modified Eagle Medium supplemented with 10% FBS, 1% penicillin-streptomycin, 1% L-glutamate, and 1% sodium pyruvate at 37℃ and 5.5% CO_2_.

### Drug treatment

Triacsin C (Enzo Biochem, Inc., BML-EI218-0100) was resuspended in DMSO at a concentration of 4.8 mM (1 mg/mL). The drug was diluted to 5 μM in complete EGM-2 for treatment. 48 hours post-infection, mock and infected cells were trypsinized and counted, then re-seeded at equivalent densities for inhibitor treatment. Q-VD-OPh (MedChem Express, HY-12305) was diluted to the indicated concentrations in complete EGM-2 alongside Triacsin C at the same time for co-treatment experiments.

### Trypan blue assay

Cell media, a PBS wash, and the trypsinized cells from each well were pooled into a single tube and centrifuged at 1000xg for 5 minutes at room temperature. Media was aspirated and cells were resuspended in a volume of media ranging from 20 μL to 100 μL, depending on the confluence of the cells at the conclusion of treatment. A 15 μL sample of the cells was taken, mixed with 15 μL of Trypan blue, and total and dead cells were counted using a hemocytometer. A minimum of 160 cells were counted per sample to ensure sufficient sample sizes.

### KSHV production and infection

BCBL-1 KSHV particles were isolated from BCBL-1 cells (5x10^5^ cells/mL) induced using 20 ng/mL of 12-*O*-tetradecanoylphorbol-13-acetate (TPA; Sigma) for 5 days as described previously. Briefly, to isolate the virions, cells were pelleted, the supernatant was passed through a 0.45 μm filter, and the virus was pelleted by centrifugation at 30,000xg for 2 hours at 4℃. The viral pellet was then resuspended in serum-free EBM-2, aliquoted, and frozen at -80℃.

To obtain KSHV.219 particles, iSLK.219 cells latently infected with KSHV.219 and various deletion mutants were selected with a combination of puromycin (1μg/mL), hygromycin B (1200 μg/mL), and G418 (250 μg/mL) for 5 days^29–31^. Following selection, iSLKs were expanded. Virus production was induced via combination treatment with doxycycline (1 μg/mL) and sodium butyrate (1 mM) for 72 hours. For the WT KSHV in Figure 3C, induction was extended to 6 to 7 days, as this batch of iSLKs did not produce as much virus. The media was then spun down at 1000 rpm for 10 min at 4℃ to remove cells. The supernatant was then centrifuged at 3000 rpm for 10 min at 4℃ to further remove any remaining cell debris. The virus was pelleted at 25,000xg for 3 hours at 4℃, resuspended in serum-free EBM-2, aliquoted, and frozen at -80℃.

Titering of these viruses was accomplished by infecting TIME cells with various dilutions of the virus stock and assessing infection success via immunofluorescence for LANA. Dilutions for experimental use were selected on the basis of yielding >85% latently infected cells and resulting in minimal background cell death. Deletion mutant viruses often required the addition of a more concentrated virus suspension to achieve similar infection rates.

5.5x10^5^ TIME cells were seeded per T25 flask, or 1.8x10^5^ TIME cells per well of a 6-well dish. 2 hours post-seeding, cells were infected with virus diluted in 1 ml of serum-free EBM-2 supplemented with polybrene at a final concentration of 8 μg/mL. The infections proceeded for 4 hours at 37℃, during which time the cells were rocked every 30 minutes. Afterwards, cells were washed with PBS and overlayed with fresh, complete EGM-2. After the establishment of latency at 48 hours post-infection, cells were used for downstream experimental applications. In each experiment, an aliquot of cells was used for LANA immunofluorescence to confirm infection rates.

### Immunofluorescence

48 hours post-infection, cells were trypsinized, and a sample of cells was seeded into a chamber slide for immunofluorescence for LANA and/or ORF59 where appropriate. Cells were given a minimum of 3 hours to sit, then were fixed using 4% PFA (m/v) in PBS for 15 min. In brief, cells were blocked using an immunofluorescence blocking buffer consisting of 1% BSA in TBS for 20 min, then probed with a primary antibody specific for LANA and a fluorophore-conjugated secondary antibody for 1 hour each with PBS washes between each step. Slides were treated with mounting media containing DAPI and imaged using a Keyence BZ-X710 fluorescence microscope.

For the assessment of p65 nuclear translocation, the same total p65 antibody was used as that used for the p65 western blots (Cell Signaling Technology Mouse Monoclonal Antibody #6956). The manufacturer’s recommended immunofluorescence protocol was followed, with the primary antibody diluted at 1:400 and incubated at 4℃ overnight, and the secondary antibody (Invitrogen A11020) diluted 1:1000 and incubated at room temperature for 2 hours. Fixation and imaging were conducted as above.

### Adenovirus production, purification, and transduction

The gutted adenoviruses used were grown in HEK 293-Cre cells as previously described^32^. In repeated rounds of infection and collection, 293-Cre cells were transduced with a pre-existing stock of AdKLAR or AdRTA and simultaneously infected with a helper adenovirus at an MOI of 1. Following each round of production, the media and cell pellet were collected and snap-frozen using liquid nitrogen, then stored at - 80℃. To generate the inoculum for the subsequent round of virus production, the pellet was snap-frozen and thawed three times to release any virus trapped in the cells, mixed with the media from the prior round, and helper adenovirus was added to achieve an MOI of 1. This inoculum was then added to cells for 1 hour at 37℃, after which the inoculum was aspirated, cells were washed with PBS, and media was replaced with complete DMEM. Media and cells were collected after the appearance of cytopathic effects at 48 hours. This was repeated and scaled up from a T25 to 10 T150s.

After the final collection, the cell pellet was snap-frozen and thawed three times to free the adenovirus within, and cell debris was removed via centrifugation at 3500 rpm for 10 min at 4℃. The supernatant from this step was loaded onto a discontinuous (15% and 40%) cesium chloride density gradient and centrifuged at 32,000 rpm for 2 hours at 4℃ in an SW41Ti rotor in a Beckman Coulter, Inc. ultracentrifuge. The mature virus band was pulled and loaded onto a second, continuous (15% to 40%) cesium chloride density gradient, and the spin was repeated. The mature virus band was pulled again, and three rounds of dialysis with A195 buffer were used to remove cesium chloride from the viral stock.

As gutted adenoviruses do not plaque, the concentration of viral particles was determined by denaturation of the viral capsid using 1.25% SDS in water, determination of DNA concentration via Qubit, an approximate MOI for optimization experiments was calculated, and expression of the KLAR was checked via RT-qPCR for LANA expression.

5.5x10^5^ TIME cells were seeded per T25 flask, allowed 2 hours to sit, and transduced with the indicated gutted adenovirus diluted at the appropriate concentration in 1 mL of serum-free EBM-2 supplemented with 1 μg/mL poly-L-lysine. The transductions were carried out for 90 minutes at 37℃, after which the cells were washed twice with PBS and provided with fresh, complete EGM-2.

### Lentivirus production and transduction

Lentiviruses were produced via co-transfection of the packaging vectors pMD2.G and psPAX2 with a payload vector encoding the gene of interest under the CMV promoter. In brief, 4.5x10^6^ HEK 293Ts were seeded in a 10 cm dish the afternoon prior to transfection, then transfected with 8 μg psPAX2, 4 μg pMD2.G, and 8 μg payload vector the following morning. Media was changed after 24 hours, and media was collected at 48- and 72-hours post-transfection, pooled, filtered through a 0.45 μm filter, and aliquoted into 1 ml aliquots.

5.5x10^5^ TIME cells were seeded per T25 flask. Cells were allowed 2 hours to sit, then transduced with 1 aliquot of lentivirus per T25, supplemented with polybrene at a final concentration of 8 μg/mL. Lentiviral transductions proceeded for 6 hours at 37℃, with cells being rocked every 30 minutes, after which the lentivirus was aspirated, cells were washed with PBS, and overlayed with fresh, complete EGM-2. 24 hours post transduction, cells were treated with puromycin (1 μg/mL) for 48 hours to select for transduced cells. After selection for the experiment shown in Figure 4D, cells were treated with Triacsin C and live cell imaged to quantify cell viability. For the lipid droplet assay, media was replaced and cells were left for 48 further hours before lipid droplet quantification, to allow lipid droplet levels in cells expressing GFP to return to levels comparable to those observed in mock infected cells (data not shown).

### Introduction of the vFLIP R12E mutation

The vFLIP R12E mutation was introduced into the WT vFLIP open reading frame via overlap extension PCR using the primers below.

vFLIP F primer: AAGCTTCGAATTCATGGCCACTTAC

vFLIP R12E R edit primer: CTGTCATCCGTGCCCAGTTTCTCCGCCACCTCACAGAGAACC

vFLIP R12E F edit primer: GGTTCTCTGTGAGGTGGCGGAGAAACTGGGCACGGATGACAG

vFLIP R primer: GACCGGTGGATCCCTATGGT

### Generation of IκBα expressing TIME cells

pBABE-Puro-IKBalpha-mut (super repressor) was obtained from Addgene (15291). The IκBα S32A/S36A open reading frame was cloned into the gateway destination vector pLenti CMV Hygro DEST via gateway cloning and both the empty vector and the resultant IκBα S32A/S36A-expressing vector were used to generate lentivirus. 24 hours post-transduction with this lentivirus, cells were placed under hygromycin selection. Hygromycin selection was maintained until mock transduced cells had died, typically after 4-6 days. Cells surviving selection were allowed to recover, and upon expansion were placed under hygromycin selection a second time, after which they were used in the indicated experiments or frozen down.

### Live cell imaging

The green, fluorescent total cell dye SYTO 24 Green (2 μM) and the red, fluorescent dead cell dye propidium iodide (2 μg/mL) were diluted in complete EGM-2. This master mix was then divided as needed for drug treatment conditions and DMSO or drug was added to achieve the desired final concentration. Cells were treated with this media, containing both dyes and DMSO or drug, allowed one hour incubation at 37℃ to allow the dyes to enter the cells, and then placed into the CellCyte live cell imager for imaging every 4-6 hours for 24-48 hours. Total and dead cell counts were assessed by the CellCyte’s analysis software, then used to calculate cell viability.

### RT-qPCR

To harvest RNA, cells were washed with PBS, then RNA was isolated using the Macherey-Nagel NucleoSpin RNA kit (740955.50). RNA was quantified via NanoDrop and 1 μg of RNA per sample was reverse transcribed using the Bio-Rad iScript Reverse Transcription Supermix for RT-qPCR (1708841). qPCR for vFLIP was conducted using the primers below, while primers for GAPDH or tubulin were used as a housekeeping normalization control. The Bio-Rad SsoAdvanced Universal SYBR Green Supermix (1725274) was used for the qPCR reactions.

vFLIP F qPCR primer: AGTGTTGGGAGTGTGATGGG

vFLIP R qPCR primer: AGAGTAGACCTACAGCGCCA

### Lipid droplet assay and flow cytometry

Nile Red (Cayman Chemical) and LipidSpot 610 (Biotium) were used with the Cayman Chemical Lipid Droplets Fluorescence Assay Kit (500001) to assess lipid droplet formation. Cells were trypsinized, pelleted at 1000xg for 5 min, resuspended in 1 mL of PBS. Cells were fixed in 500 μL of fixative solution made from the Cayman Chemical Lipid Droplets Assay Fixative (100008981) diluted 1:10 in PBS for 10 minutes, washed with 1 mL of PBS, stained with 500 μL of Cell-Based Assay Nile Red Solution (600055) diluted 1:1000 in PBS for 15 minutes or 200 μL of LipidSpot 610 diluted 1:1000 in PBS for 20 minutes, washed with 1 mL of PBS, and resuspended in a variable volume of PBS depending on the desired final cell concentration. Flow cytometry was conducted using a Guava EasyCyte for Nile Red-stained cells and using a BD FACSymphony or BD Canto RUO for LipidSpot 610-stained cells. All replicates of a given experiment were conducted using the same cytometer.

Flow cytometry data was assessed using FlowJo. In brief, forward and side scatter measurements were used to gate out dead cells and debris based on the relevant negative control cell population in the experiment. Live cell populations were then converted to histograms with the x-axis reading the relevant fluorescence on a logarithmic scale, and a range gate set using the negative control cell population was used to calculate the percentage of Nile Red+ or LipidSpot610+ live cells in each cell population. In the experiments assessing lipid droplet formation in the presence or absence of IκBα, the fluorescence range gate was set based on the cells expressing vFLIP and the empty vector.

### NF-κB Luciferase Reporter Assay

NF-κB activation by WT or mutant vFLIP was measured using the Nano-Glo Dual Luciferase Reporter Assay System (Promega). Briefly, 75,000 HEK293T cells/well were seeded into a 24 well TC-treated plate and incubated overnight at 37 oC and 5% CO2. Transfection complexes were made by incubating Mirus TransIT-LT1 reagent with a mixture of separate expression plasmids encoding KSHV vFLIP (WT or R12E) or GFP, firefly (Photinus pyralis) luciferase downstream of the NF-κB promoter, and NanoLuc luciferase containing a constitutive TK promoter to control for transfection efficiency and cell density. The ratio of NF-κB-firefly:TK-NanoLuc plasmids was 10:1 and 500 ng of total DNA was transfected into each well, in the relative amounts of 200 ng pLVX vFLIP/pLVX vFLIP R12E/pLVX GFP/pcDNA (empty), 50 ng NF-κB firefly luciferase, 5 ng TK-NanoLuc luciferase, and 245 ng pcDNA. After 40 hours of incubation, cells were lysed using 100μL of ONE-Glo EX Reagent and 100 μL of lysate was transferred to a 96-well white-walled plate. Firefly and NanoLuc luciferase levels were measured sequentially in the same sample following the manufacturer’s instructions using a CLARIOstar Plus plate reader (BMG Labtech) with an integration time of 0.8s.

To determine the relative induction of NF-κB, the compound ratio of cells transfected with either vFLIP constructs or a negative control (empty vector or GFP only plasmid) was used. First, the ratio of firefly luciferase:NanoLuc luciferase signal was determined for each well. Then, the firefly:NanoLuc ratio for vFLIP transfected cells was divided by the ratio for the negative control wells. Each data point represents the mean of technical triplicates from a single experiment and error is reported as the standard deviation of the mean across four biological replicates.

### Western blots and quantification

For the western blots assessing p65 phosphorylation, cells were transduced with the relevant lentiviruses, selected using puromycin, and collected at 120 hours post-transduction. These cells were separate populations from those collected to assess lipid droplet formation. For the blots assessing IκBα degradation, cells were collected at 120 hours post-transduction, and the cell population was split, with half of the cells being used for lipid droplet analysis and half being used for the western blots.

Cells were lysed in RIPA buffer (150 mM NaCl, 5 mM EDTA pH 8.0, 50 mM Tris-HCl pH 8.0, 1% NP-40, 0.5% sodium deoxycholate, 0.1% SDS) supplemented with Pierce Protease and Phosphatase Inhibitor EDTA-free mini tablets (Thermo) for 30 min on ice. Lysates were clarified by centrifugation at 15000g for 20 min at 4°C and concentration was determined by BCA (Thermo). Clarified lysates were denatured at 95°C for 5 minutes in 1X final concentration SDS Lamelli buffer. 5-10 ug of each sample was run on 4-12% Bis-Tris protein gels in MOPS running buffer (Invitrogen) and then transferred to Immobilon-FL PVDF membranes (Millipore Sigma) using a semi-dry Trans-Blot Turbo transfer system (BioRad). Membranes were blocked with 3% BSA in TBST and incubated overnight with primary antibody (1:1000 dilution) at 4°C. Primary antibodies used in this study are Phospho-NF-κB p65 (Ser536) (93H1) (Cell Signaling, #3033), NF-κB p65 (L8F6) (Cell Signaling, #6956), IκBα (Cell Signaling, #9242), and vinculin (Santa Cruz (7F9), sc-73614). Following 1h incubation with appropriate HRP-conjugated secondary antibodies (1:10000 dilution), blots were imaged using a Azure 600 CCD imager.

Quantification of immunoblots was performed in ImageJ. For assessing IκBα degradation, the integrated intensity corresponding to IκBα was background subtracted and normalized to background-subtacted vinculin signal in each lane. This same procedure was used to determine the induction of p65 phosphorylation with an additional step where the ratio of phosphorylated to total p65 was calculated. Finally, data were normalized to the GFP control transfection within each replicate.

### Data analysis and visualization

Data analysis and visualization were conducted using GraphPad Prism.

## Data availability

The sequences used to identify vFLIP and IκBα during the cloning process were obtained from NCBI (vFLIP gene ID: <u>4961494</u>; IκBα gene ID: <u>4792</u>). All other data from this study are included in the article itself and the supplemental material.

