## Supplementary Figures for "Kaposi sarcoma herpesvirus vFLIP induces lipid droplet formation in Tert-immortalized microvascular endothelial cells via NF-κB activation"

**Supplementary Figure 1: Measurement of expression of LANA and vFLIP**

(A) Measurement of LANA expression via RT-qPCR at 48 hours post-transduction with increasing particle:cell ratio of AdKLAR. Expression was calculated using the 2^-ΔΔCt relative to KSHV latently infected cells, with GAPDH as a loading control. (B) Measurement of LANA expression via RT-qPCR at 96 hours post-transduction with increasing particle:cell ratio of AdKLAR. Expression was calculated using the 2^-ΔΔCt relative to KSHV latently infected cells, with GAPDH as a loading control. (C) Measurement of vFLIP expression via RT-qPCR at 72 hours post-transduction and 120 hours post-transduction with vFLIP lentivirus. Expression was calculated using the 2^-ΔΔCt relative to cells expressing vFLIP at 72 hours post-transduction, with tubulin as a loading control.

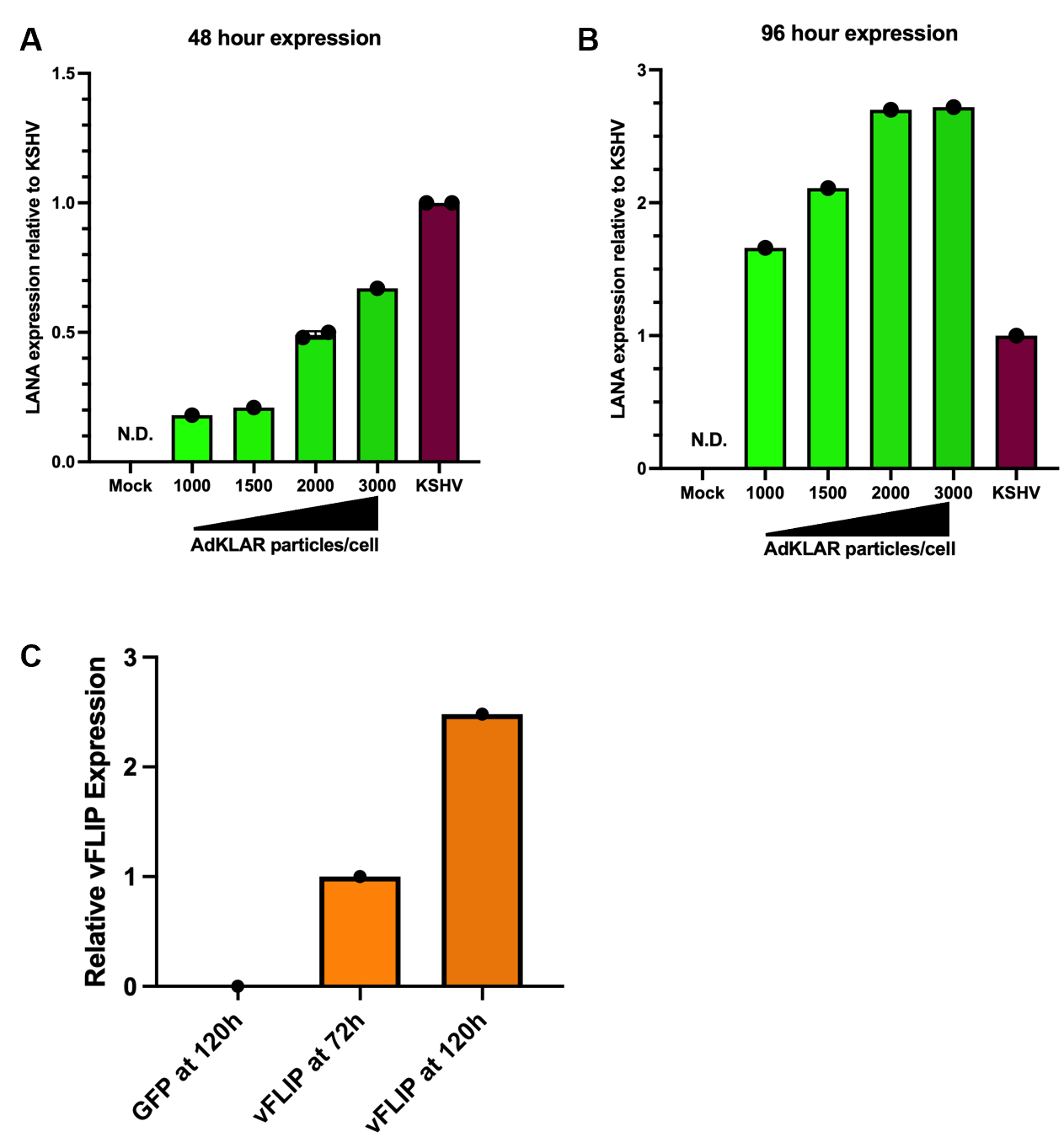

**Supplementary Figure 2: IκBα impact on lipid droplets in GFP transduced cells and on p65 phosporylation**

(A) Flow curves of unmanipulated TIME cells (red), TIME cells expressing empty vector (orange), and TIME cells expressing IκBα S32A/S36A (blue) transduced with lentivirus expressing GFP and fixed and stained with LipidSpot 610 at 120 hours post-transduction. (B) Quantification of A, with fluorescence range gate based on vector GFP cell population. Data shown are mean +/- SD, n=4, n=3 for unmanipulated TIME cells transduced with GFP. (C) Assessment of p65 phosphorylation and total p65 protein levels by western blot for vector and GFP expressing TIME cells, unmanipulated TIME cells expressing vFLIP, vector and vFLIP expressing TIME cells, and IκBα S32A/S36A and vFLIP expressing TIME cells at 120 hours post-transduction.

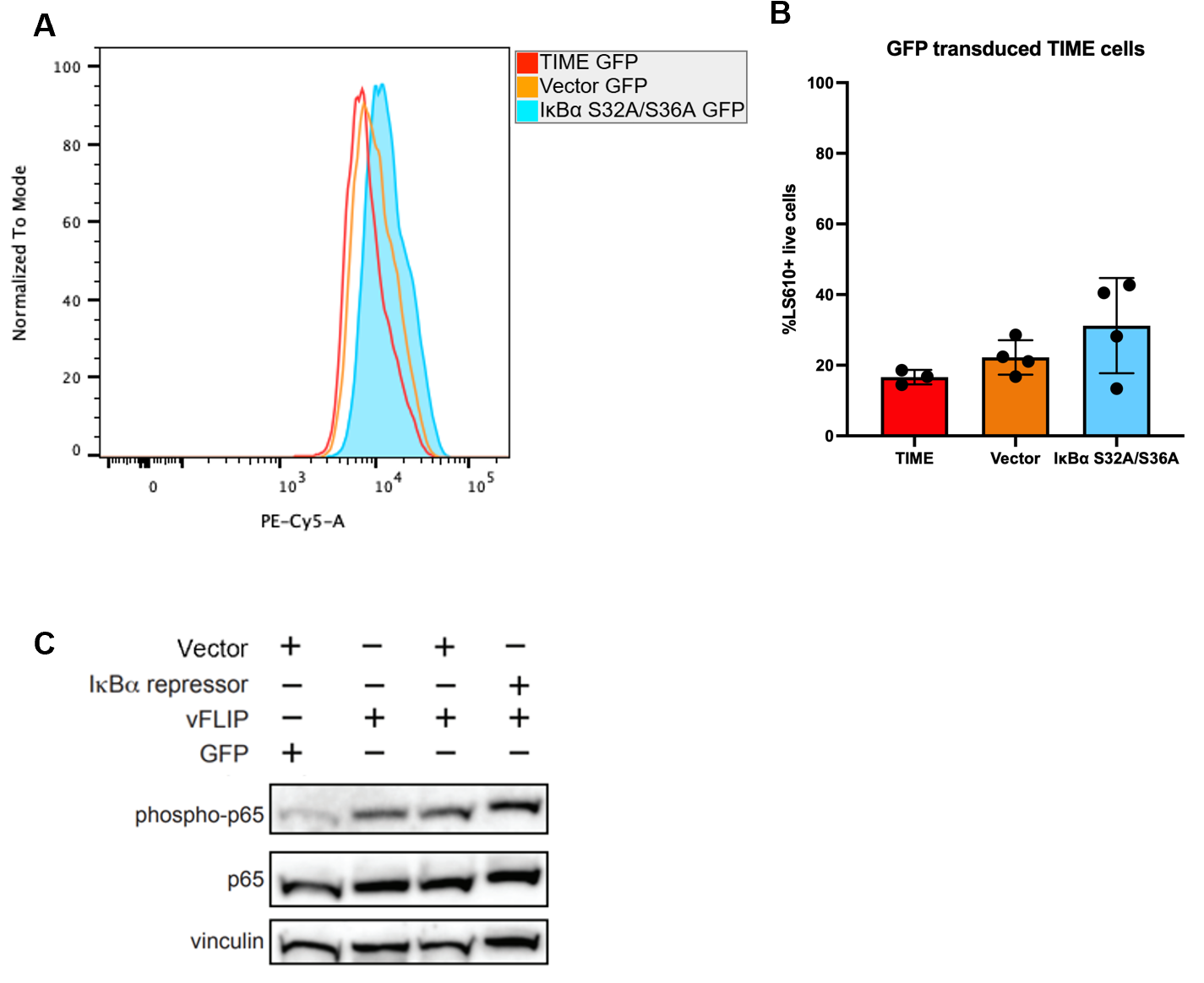
